# A novel protocol for the isolation of fungal extracellular vesicles reveals the participation of a putative scramblase in polysaccharide export and capsule construction in *Cryptococcus gattii.*

**DOI:** 10.1101/538850

**Authors:** Flavia C. G. Reis, Beatriz S. Borges, Luísa J. Jozefowicz, Bianca A. G. Sena, Ane W. A. Garcia, Lia C. Medeiros, Sharon T. Martins, Leandro Honorato, Augusto Schrank, Marilene H. Vainstein, Livia Kmetzsch, Leonardo Nimrichter, Lysangela R. Alves, Charley C. Staats, Marcio L. Rodrigues

## Abstract

Regular protocols for the isolation of fungal extracellular vesicles (EVs) are time-consuming, hard to reproduce, and produce low yields. In an attempt to improve the protocols used for EV isolation, we explored a model of vesicle production after growth of *Cryptococcus gattii* and *C. neoformans* on solid media. Nanoparticle tracking analysis in combination with transmission electron microscopy revealed that *C. gattii* and *C. neoformans* produced EVs in solid media. These results were reproduced with an acapsular mutant of *C. neoformans*, as well as with isolates of *Candida albicans, Histoplasma capsulatum,* and *Saccharomyces cerevisiae*. Cryptococcal EVs produced in solid media were biologically active and contained regular vesicular components, including the major polysaccharide glucuronoxylomannan (GXM) and RNA. Since the protocol had higher yields and was much faster than the regular methods used for the isolation of fungal EVs, we asked if it would be applicable to address fundamental questions related to cryptococcal secretion. On the basis that polysaccharide export in *Cryptococcus* requires highly organized membrane traffic culminating with EV release, we analyzed the participation of a putative scramblase (Aim25, CNBG_3981) in EV-mediated GXM export and capsule formation in *C. gattii*. EVs from a *C. gattii aim25*Δ strain differed from those obtained from wild-type (WT) cells in physical-chemical properties and cargo. In a model of surface coating of an acapsular cryptococcal strain with vesicular GXM, EVs obtained from the *aim25*Δ mutant were more efficiently used as a source of capsular polysaccharides. Lack of the Aim25 scramblase resulted in disorganized membranes and increased capsular dimensions. These results associate the description of a novel protocol for the isolation of fungal EVs with the identification of a previously unknown regulator of polysaccharide release.

**IMPORTANCE:** Extracellular vesicles (EVs) are fundamental components of the physiology of cells from all kingdoms. In pathogenic fungi, they participate in important mechanisms of transfer of antifungal resistance and virulence, as well as in immune stimulation and prion transmission. However, studies on the functions of fungal EVs are still limited by the lack of efficient methods for isolation of these compartments. In this study, we developed an alternative protocol for isolation of fungal EVs and demonstrated an application of this new methodology in the study of the physiology of the fungal pathogen *Cryptococcus gattii*. Our results describe a fast and reliable method for the study of fungal EVs and reveal the participation of scramblase, a phospholipid translocating enzyme, in secretory processes of *C. gattii*.

## Introduction

Extracellular vesicles (EVs) are produced in all domains of life (1). In fungi, these structures were first isolated in the human pathogen *Cryptococcus neoformans* (2). EVs have been further described in at least eleven additional species and their functions in fungi include the molecular transport across the cell wall (2), induction of drug resistance (3), prion transmission (4, 5), delivery of virulence factors (6, 7), immunological stimulation (8–12), RNA export (13), transfer of virulence traits (14), and trans-kingdom communication followed by regulation of expression of virulence-related genes (15). Although it is now well recognized that EVs play multiple and essential roles in fungal physiology, many questions remain unanswered (16). For instance, it is still unknown what mechanisms are required for biogenesis of EVs. The roles of EVs, if any, during infection also remain indefinite. Finally, as with other infection models, EVs have been proposed as vaccine candidates to prevent fungal diseases (8), but methods for obtaining large amounts of EVs for animal immunization are still not available. Indeed, many of the unsolved questions about fungal EVs remain unanswered because of experimental limitations. For instance, it is well-known by researchers in the fungal EV field that the standard protocols used for vesicle isolation are time-consuming (1-3 weeks) and produce very low yields (17). It is clear, therefore, that the improvement of protocols for EV isolation might solve major questions in the field.

EVs have been traditionally studied after their isolation from liquid cultures (17). However, physiological production of EVs clearly does not demand liquid matrices. For instance, EVs are now considered structural and functional components of the extracellular matrix in mammalian models (18). In this environment, they participate in matrix organization, regulation of cellular functions and determination of the physical properties of different tissues (19). EVs produced in gelatinous matrices impact tissue regeneration, inflammation and tumor progression (18, 19). With a few exceptions (e.g. blood and liquor during infection), fungal cells are distributed over solid or gelatinous matrices, including the soil, bark of trees, bird excreta and tissues of plants, insects and higher animals. Nevertheless, the production of fungal EVs in non-liquid matrices has not been explored so far, although it is reasonable to suppose that fungal cells might produce EVs in non-liquid matrices.

*C. neoformans* and *C. gattii* use fungal EVs to export virulence factors and to promote cell-to-cell communication (7, 14). Noticeably, the *Cryptococcus* model is one of the most laborious systems in which fungal EVs have been studied, due to low yields of the protocols and massive contamination with supernatant polysaccharides. EV-mediated molecular export in *Cryptococcus* demands membrane mobility (20). In this context, phospholipid flippases and scramblases are essential for membrane curvature and plasticity in different compartments of the cell (21). The flippase activity of aminophospholipid transferase 1 (Apt1) was implicated in EV production in *C. neoformans* (22, 23). The role of scramblases, however, remained unknown.

In this study we describe a novel protocol for fast and reliable isolation of fungal EVs from solid media, mostly using *C. gattii* as a model. EV isolation from solid fungal cultures revealed the participation of a putative scramblase in EV formation and surface architecture of *C. gattii*. These results reveal novel approaches and cellular regulators implicated in the study of the functions and general properties of fungal EVs.

## Results

### Fungal EVs are produced in solid media

Due to the well-known limitations of the protocols currently used for the isolation of fungal EVs from liquid media (17), we asked whether these extracellular membrane compartments would be produced in solid matrices. We hypothesized that fungal EVs could be entrapped within the fungal population grown in plates containing solid media, which would favor a relatively high density of vesicles in an area of growth limited by the plate’s dimensions. To address this question, we cultivated *C. gattii* or *C. neoformans* cells to confluence in solid YPD for 24 h, for subsequent preparation of fungal suspensions in PBS after collection of fungal cells with inoculation loops (Supplemental video 1). Cell suspensions of 30 ml at densities varying from 5 × 10^9^ to 1 × 10^10^/ml were sequentially centrifuged for removal of yeast cells and possible debris and the remaining supernatants were ultracentrifuged to collect EVs. Ultracentrifugation pellets were negatively stained and analyzed by transmission electron microscopy, which revealed the presence of vesicular structures with the typical morphology and dimensions of fungal EVs (Figure 1A). The same samples were submitted to nanoparticle tracking analysis (NTA), which revealed well-defined peaks corresponding to a major distribution of EVs within the size range of 100 to 300 nm (Figure 1B). To analyze the reproducibility of the protocol, EV isolation from *C. gattii* was independently repeated four additional times and vesicle properties were monitored by NTA. All samples produced very similar NTA profiles (Figure 1C), indicating that the protocol was reproducible. Similar preparations were analyzed by dynamic light scattering, which has been consistently used for the analysis of EV dimensions (17). The profile of size distribution was similar to that obtained by NTA (data not shown).

**Figure 1.**
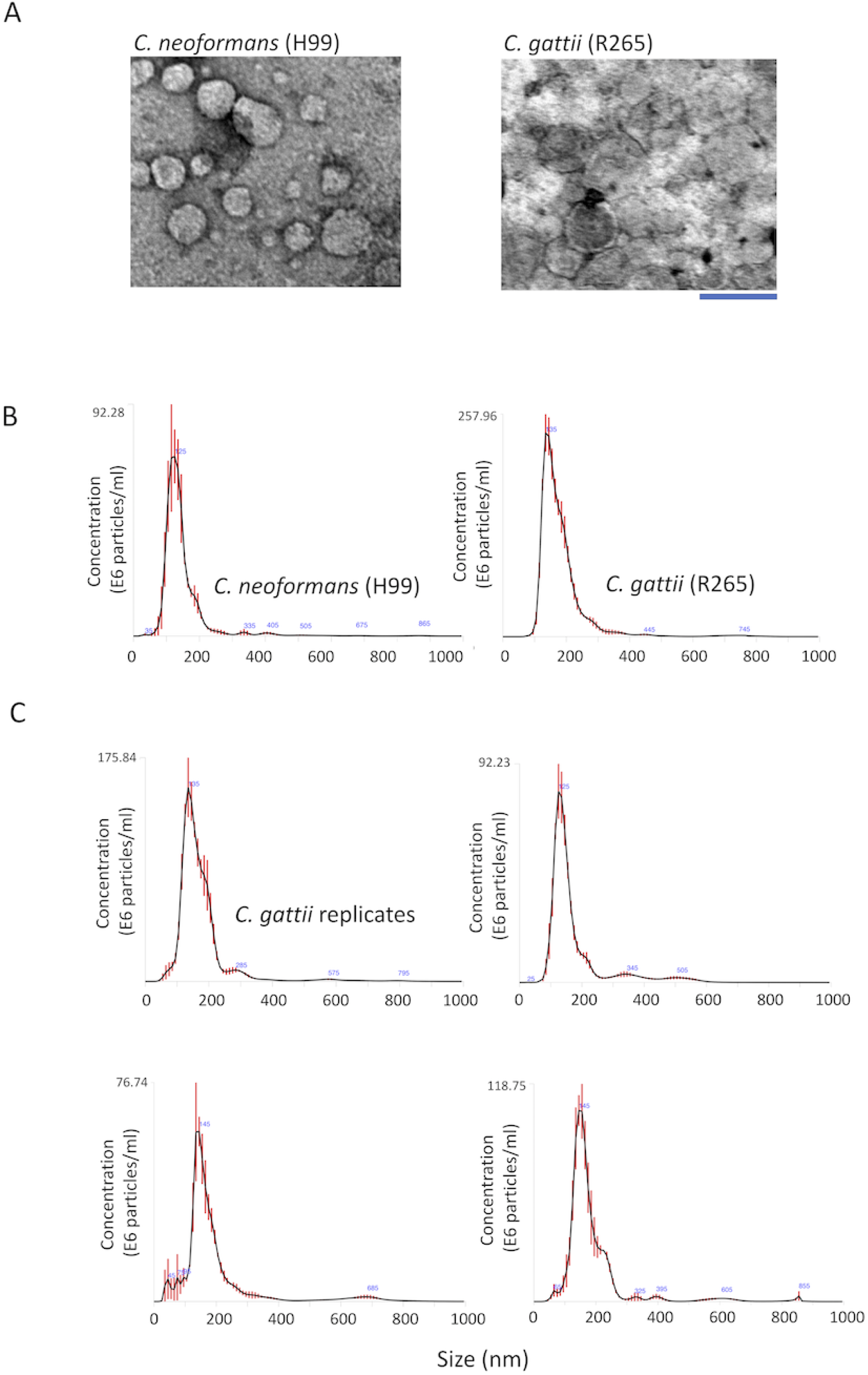
Isolation of fungal EVs from solid cultures of *C. neoformans* and *C. gattii* (strains H99 and R265, respectively). A. Transmission electron microscopy of vesicular fractions obtained after growth of both pathogens on solid YPD. Scale bar, 200 nm. B. Nanoparticle tracking analysis of the EV preparations illustrated in A, showing a concentration of vesicles in the range of 100 to 200 nm. C. NTA profiles of four samples of *C. gattii* EVs obtained independently.

We then asked if EV detection in solid media would only occur under specific experimental conditions. To address this question, we analyzed the production of EVs in a different medium or using distinct fungal species or strains. We first checked whether *C. gattii* and *C. neoformans* produced EVs in Sabouraud’s medium. Vesicular structures were abundantly detected, but the profile of size distribution included a minor population ranging from 300 to 600 nm in size (Figure 2A). Since cryptococcal EVs have long been associated with the export of GXM (2), we also asked whether EV detection after growth in solid media would be influenced by the presence of the capsule. We then analyzed vesicles obtained from an acapsular mutant of *C. neoformans*. As revealed by NTA, the *cap67*Δ strain of *C. neoformans* also produced EVs (Figure 2B). To investigate whether EV detection in solid media is exclusive to the *Cryptococcus* genus, we analyzed EV samples produced by *C. albicans, S. cerevisiae and H. capsulatum*. In all cultures tested, NTA revealed particles with properties that were compatible with EVs in size distribution. However, while *C. albicans* and *S. cerevisiae* gave nanoparticle signals that were similar to those found in *C. neoformans* and *C. gattii, H. capsulatum* produced EVs with a more diverse size distribution (Figure 2C). Together, these results indicate that EV production in solid media is a general and consistently reproducible phenomenon.

**Figure 2.**
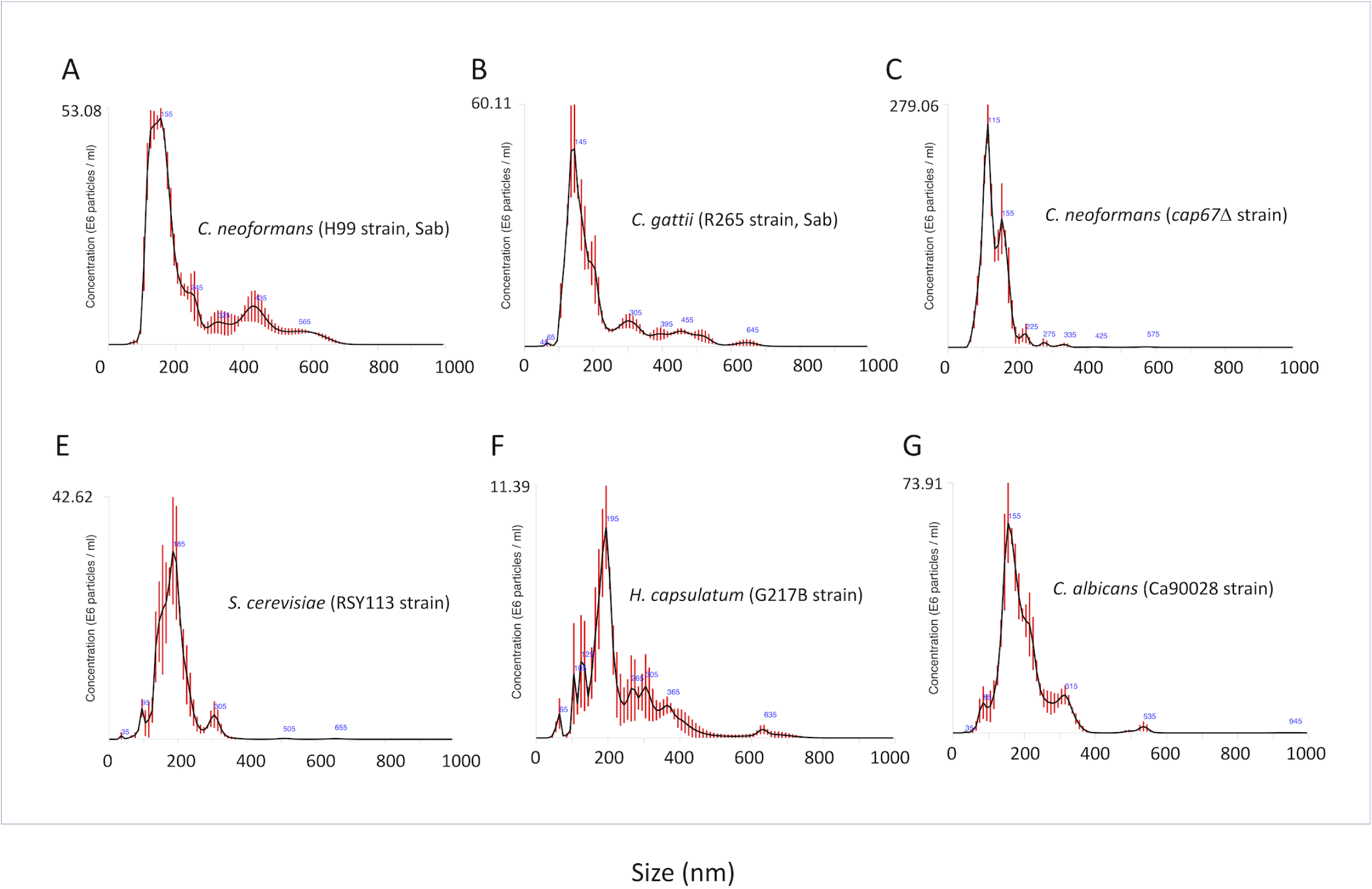
NTA profiles of EVs obtained from different fungal cultures in solid media. A-B. Analysis of *C. neoformans* (A) and *C. gattii* (B) EVs obtained from Sabouraud cultures. C. NTA of EVs obtained from an acapsular mutant of *C. neoformans*, suggesting that vesicle release in solid medium does not demand capsular structures. EVs were also detected by NTA after growth of *S. cerevisiae* (E), *H. capsulatum* (F) and *C. albicans* (G), indicating that the protocol is applicable to the study of different fungal pathogens.

### RNA and GXM are components of cryptococcal EVs produced in solid media

Following the detection of EVs in solid cultures of fungal cells, we asked whether the cryptococcal membrane particles would contain the typical components that were previously described in fungal samples of EVs (2, 24). Since different RNA classes were previously characterized as components of EVs produced by *C. neoformans, Malassezia sympodialis, Paracoccidiodes brasiliensis, C. albicans*, and *Saccharomyces cerevisiae* (13, 25), we investigated whether these nucleic acids were present in vesicle samples obtained from solid cultures of *C. gattii* and *C. neoformans*. Bioanalyzer analysis revealed the presence of RNA in EVs produced by both species (Figure 3). The RNA pattern was similar to that observed for other eukaryotes and most of the RNA was composed of molecules smaller than 200 nt, with a peak around 20-25 nt. The profiles of RNA detection were similar in both species (Figure 3A).

**Figure 3.**
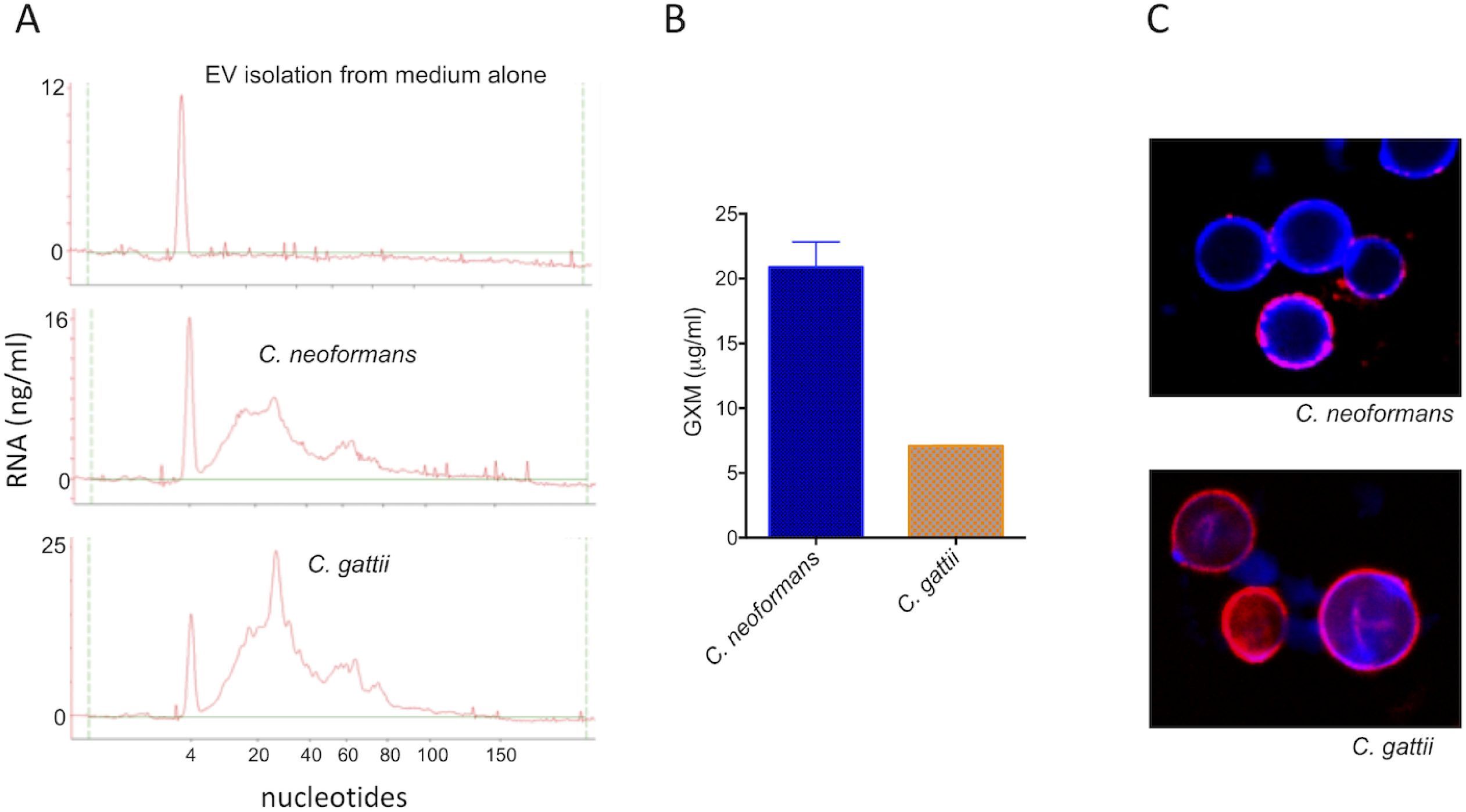
Analysis of the cargo of fungal EVs obtained from solid cultures of *C. neoformans* and *C. gattii* (strains H99 and R265, respectively). A. Analysis of nucleic acid content confirmed the presence of small RNAs in cryptococcal vesicles. In these panels, the Y-axis corresponds to RNA detection as a function of fluorescence intensity, while the X-axis represents RNA size in nucleotides. The first, sharp peak at 4 nucleotides corresponds to the RNA size the marker. No RNA was detected in control samples obtained from the culture medium. B. Detection of GXM in vesicular samples obtained from *C. neoformans* and *C. gattii* by ELISA. The concentration of vesicular GXM was significantly higher in *C. neoformans* samples (*P* = 0.0099). C. Functional analysis of vesicular GXM in samples obtained from solid medium. The *cap67*Δ mutant of *C. neoformans* efficiently incorporated GXM (red fluorescence) from EVs produced by both *C. neoformans* and *C. gattii* in solid medium into the cell wall (blue fluorescence). Results in all panels are representative of three independent experiments.

GXM is another major component of cryptococcal EVs (2). We then analyzed *C. neoformans* and *C. gattii* samples for the presence of this polysaccharide by ELISA. EVs were disrupted by treatment with organic solvents and GXM-containing precipitates were tested for reactivity with a monoclonal antibody to GXM (mAb 18B7), which confirmed the presence of the polysaccharide (Figure 3B). The results differed in *C. neoformans* and *C. gattii*, with the latter showing a significantly lower amount of vesicular GXM. These vesicular GXM samples were used in assays of polysaccharide incorporation into the surface of the *cap67*Δ acapsular mutant of *C. neoformans*. These cells efficiently incorporated the vesicular polysaccharide into their cell surface (Figure 3C).

### A putative scramblase participates in EV-mediated export in *C. gattii*

On the basis of the consistent detection of EVs in solid cultures of *C. neoformans* and *C. gattii*, we asked whether the protocol would be applicable to address biological questions related to fungal secretion. Since scramblases are directly linked to membrane traffic and secretion (21), we selected a putative scramblase that has been previously suggested in the *C. gattii* model as a regulator of secretion and target for antifungals (26) as a possible regulator of EV formation and/or polysaccharide release. The gene encoding the putative scramblase (*AIM25*, CNBG_3981) was knocked down in the R265 background of *C. gattii* and the resulting mutant cells were phenotypically characterized. The mutant had normal proliferation rates (not shown) and produced EVs in solid medium (Figure 4A). The amounts of EVs produced by mutant cells tended to be smaller, but no statistical significance was observed (data not shown). Deletion of *AIM25*, however, resulted in the production of a population of EVs of higher dimensions, in comparison to vesicles produced by wild-type *C. gattii* (Figure 4B). Vesicular components were also affected in the *aim25*Δ mutant, as concluded from the altered profile of RNA detection in mutant EVs (Figure 4C).

**Figure 4.**
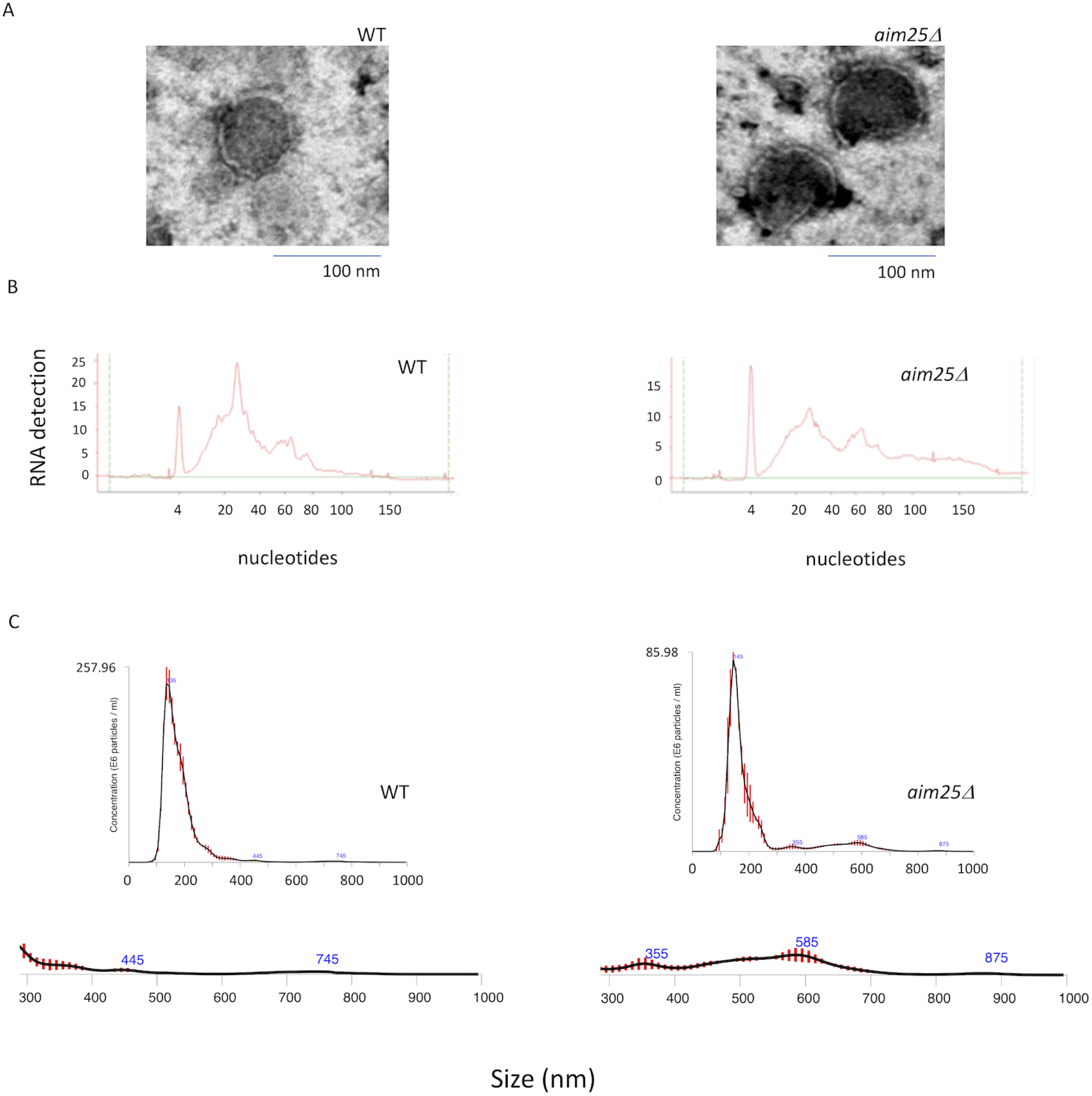
Analysis of EVs obtained after growth of wild-type (WT) or mutant (*aim25Δ*) cells of *C. gattii* in solid medium. A. Transmission electron microscopy of WT and mutant cells. B. Analysis of small RNAs contained in EVs produced by WT and mutant cells. C. NTA of EVs produced by WT and mutant cells, suggesting an increased detection of larger EVs (300-900 nm) in mutant cultures. The 300-900 nm size range of EVs was amplified in the bottom of panel C. Results shown in panels B and C are representative of two and three independent experiments, respectively.

The differences in EV dimensions and cargo were suggestive of a role of the putative scramblase in membrane organization and / or EV biogenesis. In fact, transmission electron microscopy (TEM) revealed that *C. gattii aim25*Δ mutant cells had clearly disorganized membranes, which included a general lack of the typical cryptococcal vacuoles, aberrant membranous structures, atypical plasma membrane invaginations and linearized membranous filaments with no apparent connections with cellular organelles (Figure 5). These results were consistent with the primary roles played by scramblases in the membrane organization of other eukaryotic cells (21).

**Figure 5.**
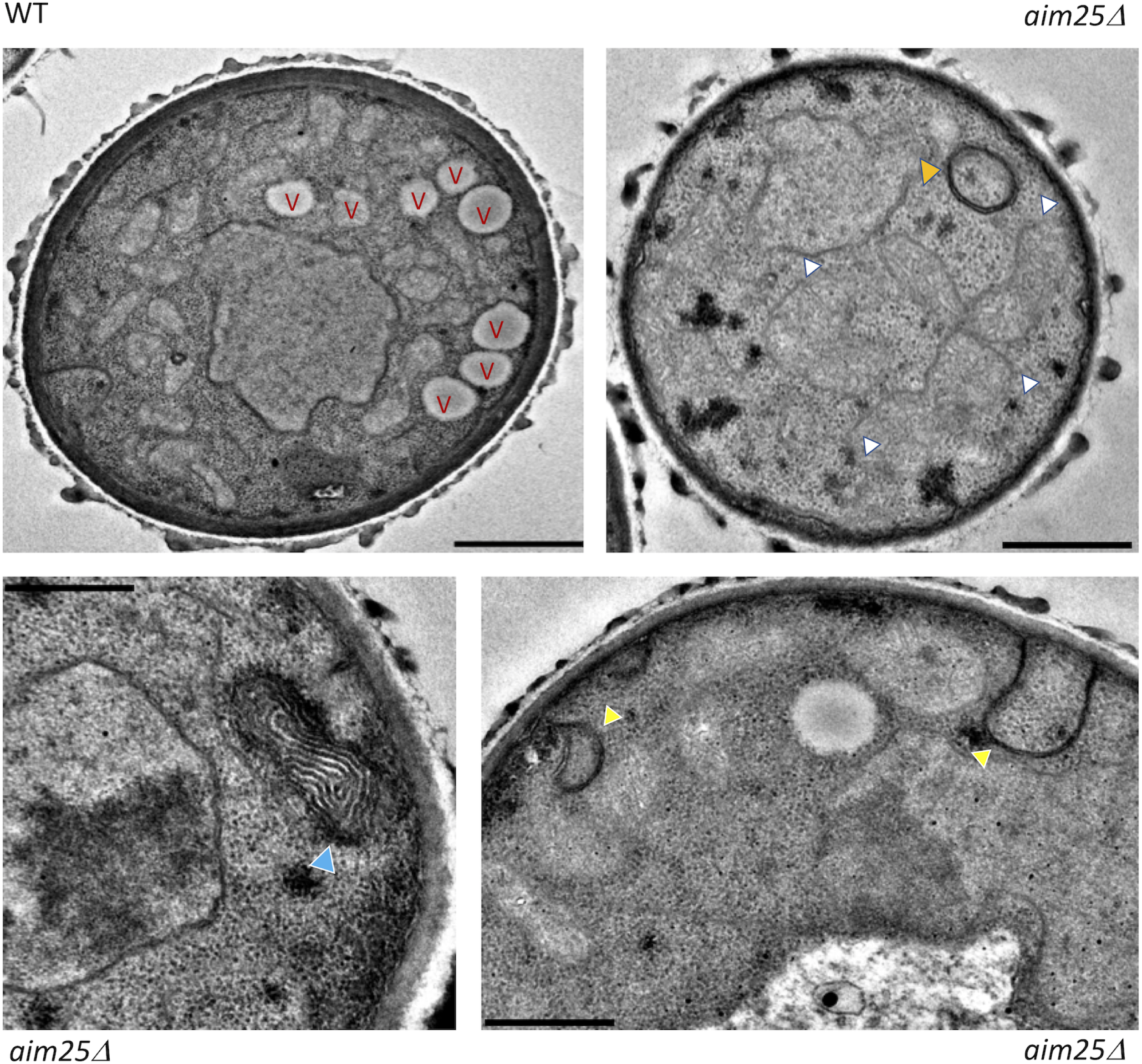
Transmission electron microscopy of wild-type (WT) and mutant (*aim25Δ*) cells of *C. gattii*. WT cells manifested the typical intracellular morphology of cryptococci, including well-defined vacuoles (V) and organized membranous compartments. In mutant cells, distorted membranes were abundantly detected. Phenotypic traits that were exclusive to mutant cells included a general lack of the typical cryptococcal vacuoles, highly electron dense membranous compartments (orange arrowhead), linearized membranes (white arrowheads), electron dense, stacked membranes (blue arrowhead) and atypical invaginations of the plasma membrane (yellow arrowheads). Scale bars correspond to 200 nm.

Polysaccharide export in *Cryptococcus* relies on membrane mobility and vesicular traffic (2, 20). Therefore, we quantified GXM in vesicular and non-vesicular extracellular fractions of wild-type and mutant cells of *C. gattii*. In comparison with parental cells, the concentration of GXM was much higher in the supernatants of the *aim25*Δ mutant strain (Figure 6A). However, no significant differences were observed when GXM was quantified in vesicular fractions, although the mutant tended to produce reduced amounts of EV-associated GXM. We, therefore, hypothesized that in the absence of scramblase, GXM could be more efficiently released from EVs, becoming more abundant in soluble supernatant fractions. To address this question, we compared the ability of acapsular *cap67*Δ cells to incorporate vesicle-associated polysaccharide obtained from wild-type cells and the *aim25*Δ mutant. After 24 h of incubation, the acapsular *cap67*Δ cells incorporated GXM from EVs produced by the *aim25*Δ scramblase mutant more efficiently than from wild-type vesicles, as concluded by immunofluorescence (Figure 6B) and flow cytometry (Figure 6C) analyses. This result agrees with a more efficient extraction of GXM from vesicles obtained by fungal cells lacking *AIM25*.

**Figure 6.**
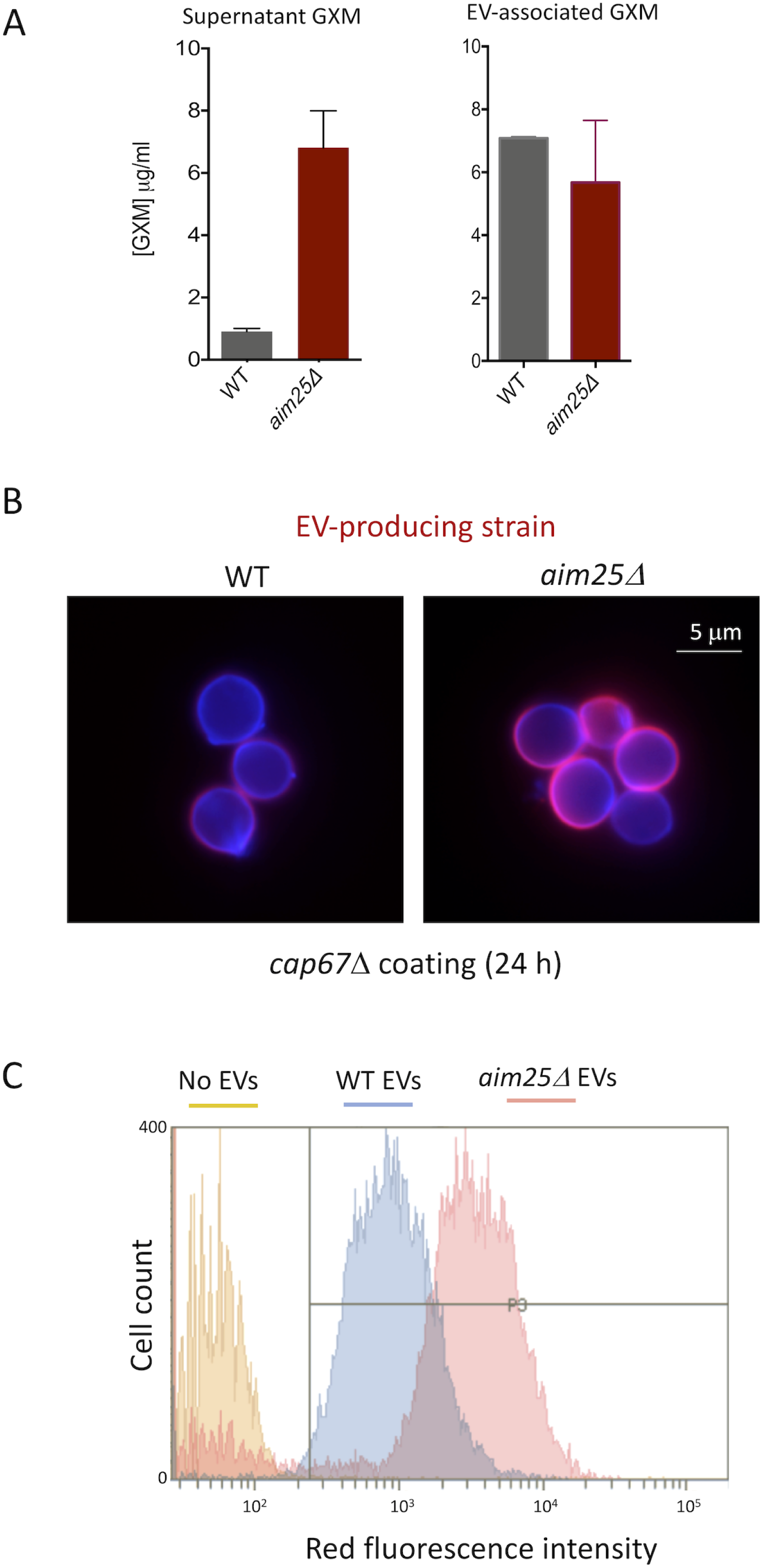
Analysis of extracellular GXM in wild-type (WT) and mutant (*aim25Δ*) cells of *C. gattii*. A. Determination of extracellular GXM in supernatant samples (A) demonstrated that mutant cells produced significantly increased polysaccharide concentrations (*P* = 0.0001). No significant changes in the GXM content (*P* = 0.41) were observed in EV samples. B. Microscopic examination of the ability of *cap67Δ* cells to incorporate GXM obtained from *C. gattii* suggested that GXM incorporation by the acapsular strain was more efficient when *aim25Δ* vesicles were used. Blue fluorescence denotes cell wall staining with calcofluor white. Red fluorescence corresponds to GXM staining with mAb 18B7. C. Flow cytometry analysis of acapsular cells under the conditions described in B, providing a quantitative confirmation of the visual observation resulting from microscopic analysis. Results are representative of two independent experiments.

Since GXM export in EVs and increased concentration of capsular polysaccharides in supernatants were linked to capsular enlargement (2), we asked if the *aim25*Δ mutant was more efficient in producing large capsules than wild-type cells of *C. gattii*. Wild-type and mutant cells had their capsular morphology analyzed by scanning electron microscopy. We first analyzed fungal cells under the conditions of EV isolation. Growth in YPD inhibits capsule formation (27) and, as expected, capsular dimensions were reduced in both wild-type and *aim25*Δ mutant cells cultivated in the solid medium (Figure 7). A closer analysis of fungal cells, however, indicated that capsule fibers, although small in dimensions, were more abundant in mutant cells grown in YPD (Figure 7A-D). We then asked whether eventual differences in capsular structures would become more evident under conditions of capsule induction in RMPI (27). This approach in fact resulted in fungal cells with larger capsules (Figure 7E-G). Under these conditions, yeast cells with more exuberant capsules were apparently more frequently observed in *aim25*Δ mutant populations. To confirm the visual perception that capsule enlargement was facilitated in mutant cells, we quantified the average capsular dimensions in conditions of capsule repression (YPD) and induction (RPMI) (Figure 8). Despite the increased number of capsular fibers in mutant cells, no differences in capsular dimensions were observed after growth of *C. gattii* in YPD. Capsule induction in RPMI, however, was significantly more efficient in mutant cells. Together, these results indicate that deletion of *AIM25* resulted in a more efficient extracellular release of GXM resulting in facilitated capsule enlargement.

**Figure 7.**
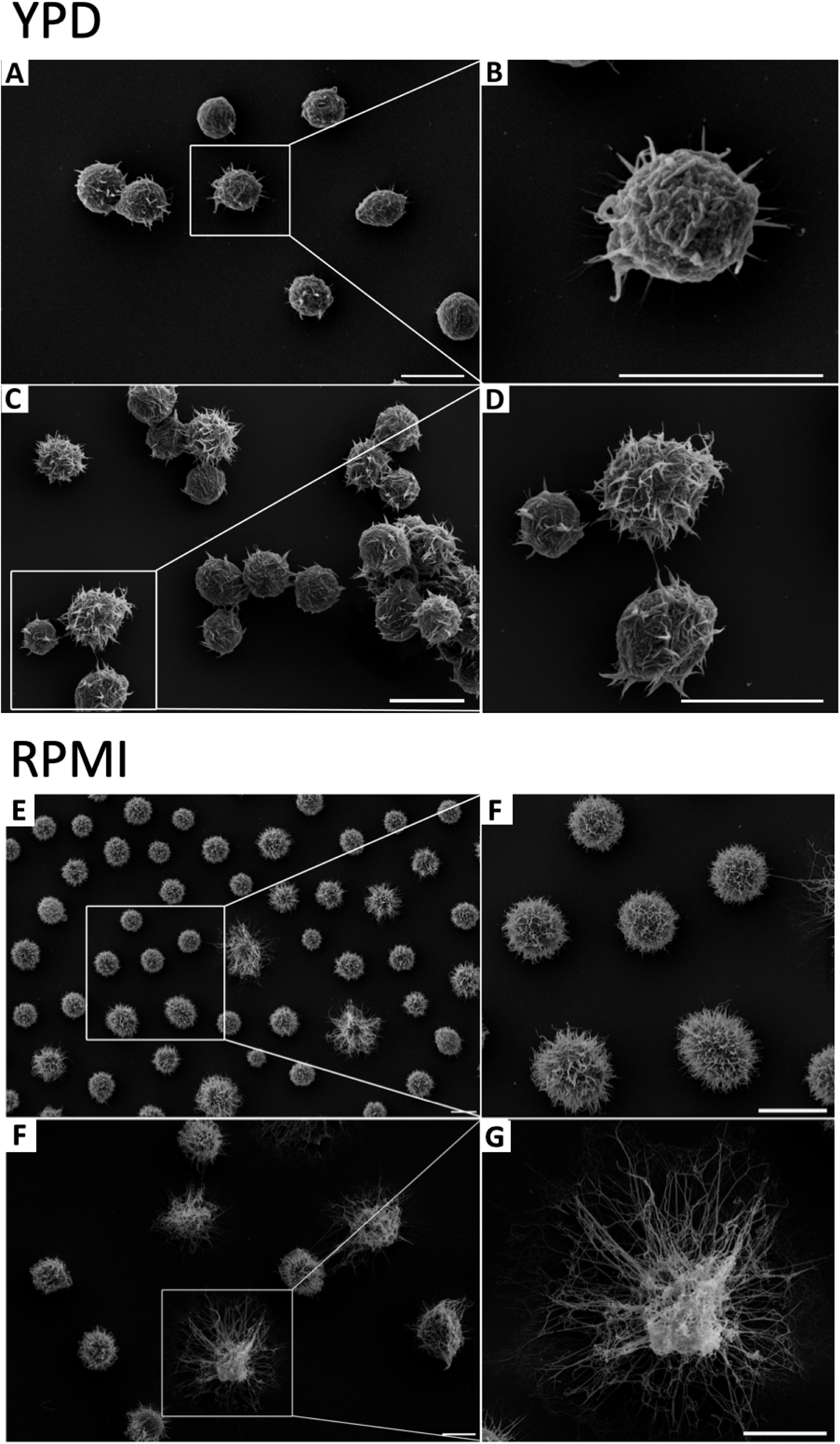
Scanning electron microscopy of wild-type (WT) and mutant (*aim25Δ*) cells of *C. gattii* after growth in solid YPD (capsule repression) or incubation in RPMI (capsule induction). General views of WT (A, E) or *aim25Δ* (C, F) cells are shown for each condition in the left panels. Magnified views of WT (B, F) or *aim25Δ* (D, G) cells are shown in the right panels. Scale bars correspond to 5 μm.

**Figure 8.**
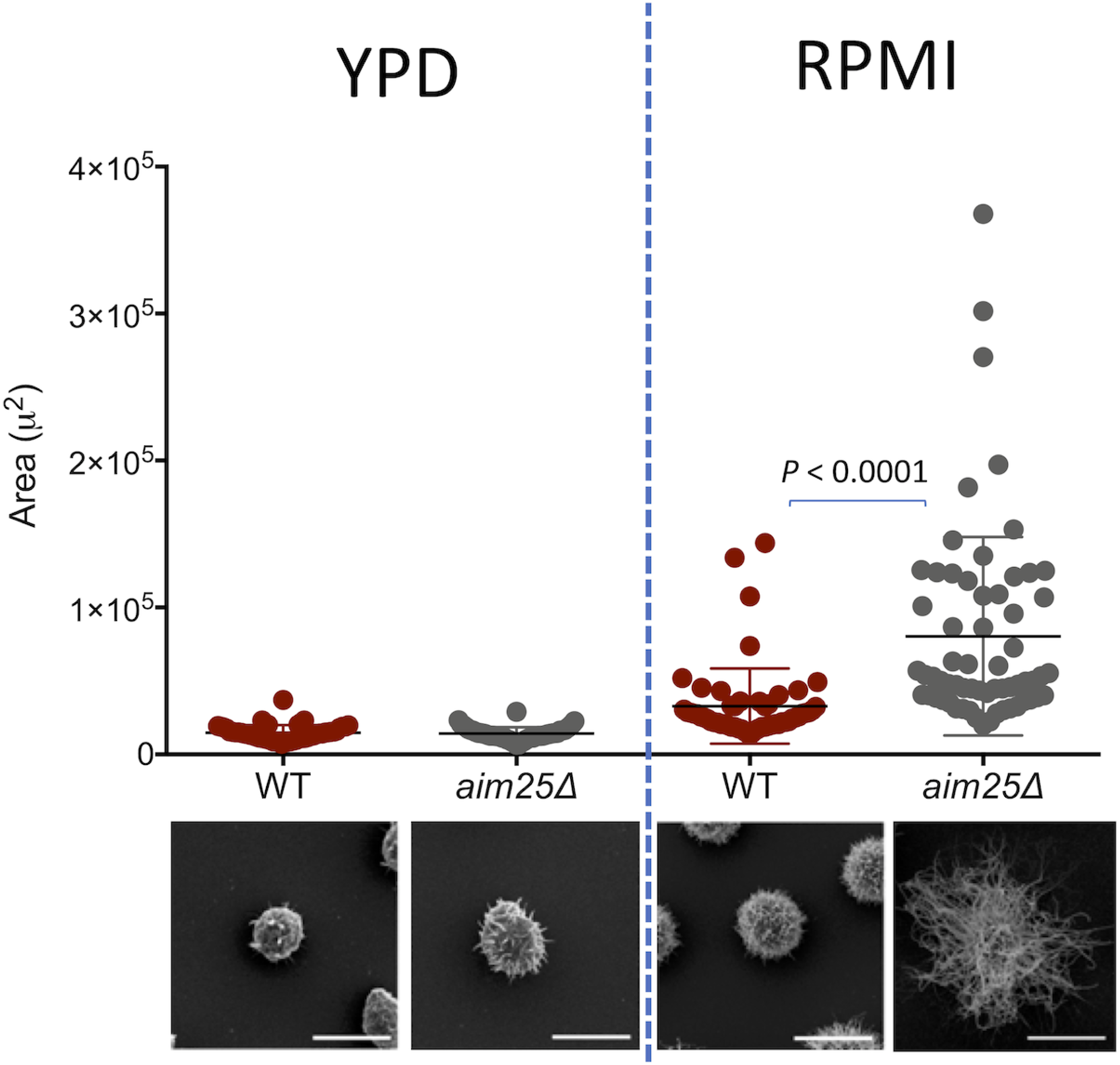
Analysis of cellular area as a consequence of capsular dimensions in wild-type (WT) and mutant (*aim25Δ*) cells of *C. gattii* after growth in solid YPD (capsule repression) or incubation in RPMI (capsule induction). The most representative phenotypes observed in each experimental condition are shown on the bottom as scanning electron microscopy illustrations. Differences in capsular dimensions had no statistical significance, with the exception of the comparison between WT and mutant cells after incubation in RMPI. Scale bars correspond to 5 μm.

## Discussion

Fungal EVs have been implicated in a number of biological processes, including transfer of virulence and antimicrobial resistance (3, 14). Neutralization of EV-mediated export has been suggested as a therapeutic strategy (15). However, several aspects of the biology of fungal EVs remain to be elucidated (16). The reduced knowledge on the characteristics of fungal EVs is likely a consequence of inefficient protocols, which usually involve centrifugation of liters of cultures, massive loss of biologically active samples and very low yields (17). In addition, there are no efficient protocols for gradient-based separation of fungal EVs. Consequently, compositional analysis of different populations of fungal EVs is highly complex, which impacts studies on their biogenesis and immunological functions (28). Therefore, although fungal EVs have been proposed as regulators of virulence, morphological transitions and immunity [reviewed in (29)], it remains elusive if they are produced during infection or if they are produced by filamentous forms of fungi. The bioactive EV components mediating antifungal resistance and virulence transfer are also unknown. Finally, it is still not clear if fungal EVs correspond to exosomes, microvesicles, cytoplasmic subtractions or a combination of all of them (30). The development of more efficient protocols is, therefore, necessary for studies aiming at understanding composition and biological functions of fungal EVs of different nature.

Fungal pathogens are rarely found in liquid matrices. Excepting the cases of fungemia and liquor contamination, pathogenic fungi are usually colonizing tissues, mucosae and the extracellular matrix during infection (31). In the environment, fungal species with pathogenic potential to humans and animals are usually distributed into the soil, tree shelves and animal excreta (31). On the basis of this assumption, we hypothesized that fungal EVs could be produced in solid matrices. Besides the biological aspects of EV production in solid media, isolation of vesicle preparations would be facilitated by easier control of both area and volume limitation imposed by flasks or plates containing solid medium. We therefore designed a protocol through which extracellular fungal components would be collected from solid media and suspended in relatively reduced volumes for further ultracentrifugation. This protocol was efficient for different fungal species, highly reproducible and essentially fast. EVs isolated from solid media were shown to be biologically active and to contain at least some of the typical components of EVs, as concluded from experiments demonstrating the traffic of GXM and presence of RNA, respectively. Importantly, while the protocols available for isolation of fungal EVs could take up to weeks and include numerous rounds of supernatant concentration and ultracentrifugation (32), the proposed protocol took approximately 5 h from collection of extracellular components to NTA analysis without any additional cost. We also showed that the facilitated protocol for EV isolation was applicable to address important biological questions related to fungal export. For example, our study compared for the first time the properties of EVs produced by *C. neoformans* and *C. gattii*, which revealed important differences in polysaccharide content.

We extended the currently described approach to the study of potential regulators of EV-mediated export. Scramblases and flippases are different types of enzymatic groups of phospholipid transportation enzymes (21). It is reasonable to suppose that regulators of membrane architecture are required for proper EV release. Indeed, in *C. neoformans*, the Apt1 flippase regulated EV physical properties and GXM export (22, 23). The participation of other membrane regulators remained unexplored, as well as the role of phospholipid translocators in the *C. gattii* model. In this context, we selected a putative scramblase (Aim25) as a candidate of regulator of membrane architecture and EV formation in the *C. gattiii* model.

Both WT and *aim25*Δ mutant cells lacking the gene putatively encoding the *C. gattii* scramblase produced EVs. Based on the detection of a population of larger EVs in cultures of mutant cells, we hypothesized that scramblase was required for membrane organization, proper EV formation, and extracellular cargo release from fungal cells. Membrane organization and proper EV formation were in fact affected in *aim25*Δ mutant cells, as concluded from TEM and NTA analyses. EV cargo was likely impacted in mutant cells, as concluded from the altered profile of RNA detection in mutant vesicles. Unexpectedly, GXM concentration was highly increased in whole supernatants of *aim25*Δ mutant cells. We therefore hypothesized that the altered membrane organization of mutant cells could result in a more efficient release of GXM from EVs. If this hypothesis was valid, a more efficient capsule formation would be expected in *aim25*Δ mutant cells. Our results demonstrated that *cap67*Δ acapsular cells were indeed more efficient in uptaking GXM from EVs produced by the *aim25*Δ mutant. In fact, these scramblase mutant cells had more exuberant capsules. These results contradict the notion that deletion of membrane regulators with negatively impact GXM export and capsule formation and efficiently illustrate the complexity of the physiological functions of EV-mediated molecular export.

The impact of scramblase deletion in *Cryptococcus* is likely broader. For instance, mutant cells of *C. neoformans* lacking flippase expression had aberrant Golgi structure, attenuated synthesis of phospholipids, increased production of immunogenic sterols and reduced formation of virulence-related lipids (22, 23). Flippase and scramblase functions, however, are not necessarily related. In contrast to our current observations, flippase mutants had decreased GXM synthesis and reduced capsular dimensions (22).

Our present results describe a novel, simplified protocol of EV isolation and its application to reveal functions of a previously unknown regulator (Aim25 scramblase) of EV properties in *Cryptococcus*. The impact of the use of the new approach to study fungal EVs will be revealed in the future, but considering the well-known difficulties in the field, it is expected that the protocol will be useful not only to identify other regulators of EV formation, but also to address the immunological functions of fungal EVs and to develop new alternatives to the study of their biogenesis and composition. The potential of this methodology to investigate EV formation in different fungal species and morphological stages is also foreseeable.

## Material and methods

### Fungal strains

The isolates used in this study included the standard strains H99 of *C. neoformans* and R265 of *C. gattii*, the *C. albicans* strain 90028, the *S. cerevisiae* strain RSY113 and the *H. capsulatum* strain G217B. The acapsular mutant *cap67Δ* of *C. neoformans* was used for glycan incorporation assays. The Delsgate methodology was used to construct the *Δaim25* (CNBG_3981) mutant strain lacking scramblase expression. Two fragments (∼ 1000 bp) encompassing the 5’ and 3’ flanking sequences of the CNBG_3981 locus were PCR amplified and gel purified using the PureLink Quick Gel Extraction & PCR Purification Combo Kit (Invitrogen). Both fragments were mixed (100 ng of each) with pDONR-NAT vector (∼200 ng), as previously described (33) and submitted to BP clonase reaction, according to manufacturer’s instructions (Invitrogen). The cassette was transformed in *Escherichia coli* OMINIMAX cells and selected by antibiotic resistance screening and colony PCR. Biolistic transformation was performed to introduce the deletion construct previously linearized by I-SceI enzymatic digestion in *C. gattii*, as previously described (33). The screening was performed using nourseothricin resistance and colony PCR. The mutant strain was confirmed by semi-quantitative RT-PCR using actin transcripts as a loading control, according to protocols previously used (33). Primers used are listed in Table 1. Stock cultures of *C. neoformans, C. gattii, S. cerevisiae* and *C. albicans* were maintained through passages in Sabouraud plates. H. capsulatum was kept in brain heart infusion agar supplemented with sheep blood (5%).

**Table 1:**
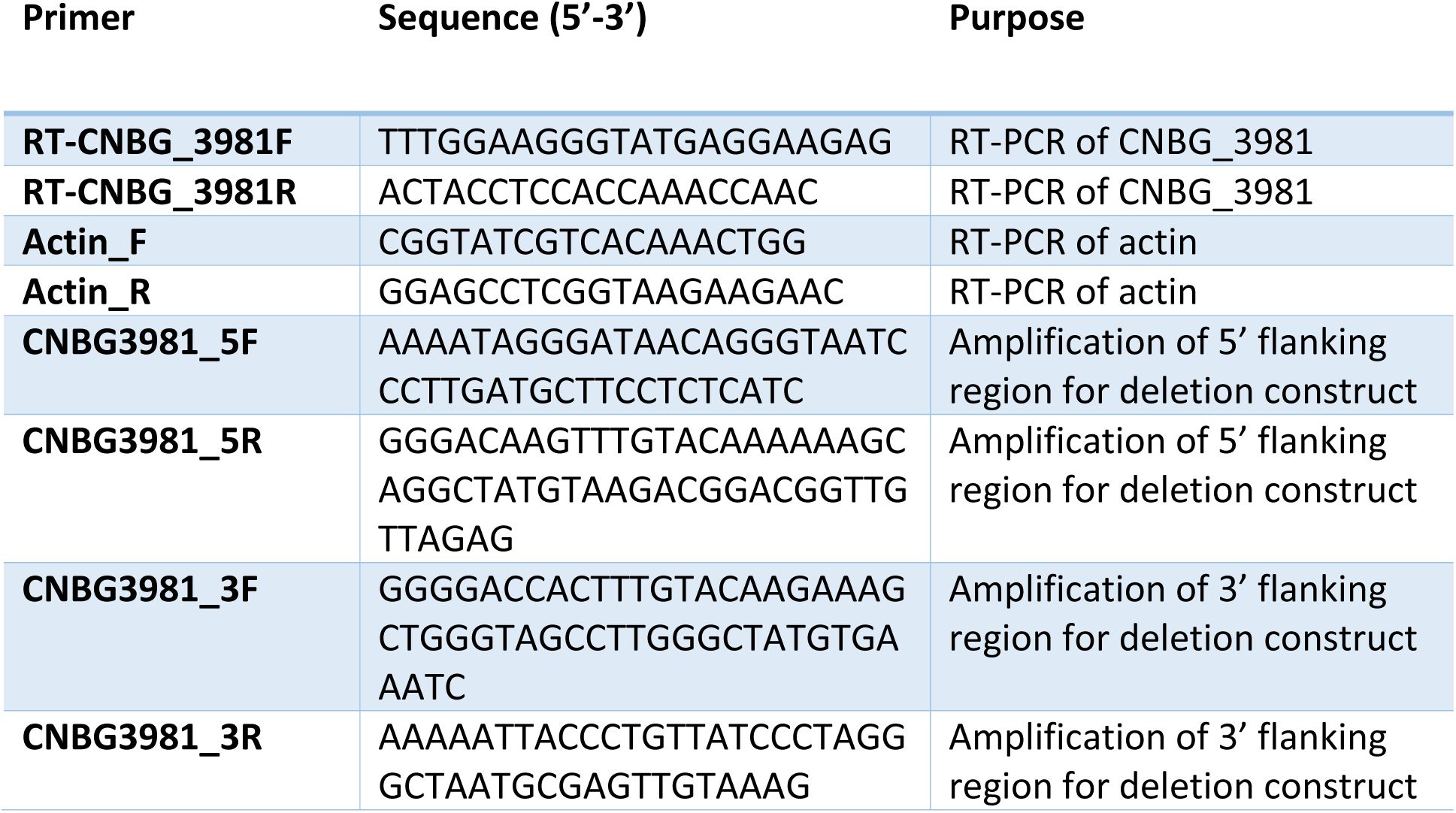
primers used for deletion of a putative scramblase (CNBG_3981, Aim25) of *C. gattii*.

### EV isolation from solid media

One colony of each isolate cultivated in solid Sabouraud was inoculated into 5 ml of yeast peptone dextrose (YPD) media and cultivated for 2 days at 30°C, with shaking. Due to specific nutritional requirements, the only exception was *H. capsulatum*, which was cultivated in Ham’s F12 medium for 48 h at 37°C, with shaking. The cells were counted and diluted to the density of 3.5 × 10^7^cells / ml in YPD. Aliquots of 300 μl of these cell suspensions were spread into YPD-agar plates (90 × 15 mm petri dishes containing 25 ml of medium) and incubated for 1 day at 30°C to reach confluence. Once again, the exception was *H. capsulatum*, which was cultivated in Ham’s F12 agar and incubated for 48 h at 37°C. Three petri dishes were used for each EV isolation. The cells were gently recovered from each of the three plates with an inoculation loop and transferred to a single centrifuge tube containing 30 ml of PBS (Supplemental video 1) previously sterilized by filtration through 0.22 μm membranes. Suspended cells were collected by centrifugation at 5,000 *g* for 15 min at 4°C. The supernatants were collected and centrifuged again at 15,000 *g* for 15 min at 4°C, to remove debris. The resulting supernatants were filtered through 0.45 μm syringe filters and centrifuged at 100,000 *g* for 1 h at 4°C. Supernatants were discarded and pellets suspended in 300 μl of sterile PBS. EV preparations were maintained at 4°C. The presence of EVs was monitored by nanoparticle tracking analysis and electron microscopy, as detailed below.

### Transmission electron microscopy

Fungal cells were washed twice in PBS and fixed for 1 h in 2.5% glutaraldehyde in 0.1 M phosphate buffer at room temperature. The fixed yeast cells were washed twice in 0.1 M cacodylate buffer and then post-fixed with 1% osmium tetroxide / 1.6% potassium ferrocyanide / 5 mM CaCl_2_ diluted in 0.1 M cacodylate buffer for 30 min at room temperature. The samples were washed three times with 0.1 M cacodylate buffer, dehydrated in a graded acetone series (5 min in 30, 50, 70, 90 and 100%) and then embedded in PolyBed812 resin. Ultrathin sections were obtained in a Leica EM UC6 ultramicrotome, collected on copper grids, contrasted with 5% uranyl acetate and lead citrate and then visualized in a JEOL 1400Plus transmission electron microscope at 90 kV. For negative-stain electron microscopy of EVs, samples obtained from solid media were transferred to carbon- and Formvar-coated grids and negatively stained with 1 % uranyl acetate for 10 min. The grids were then blotted dry before immediately observing in a JEOL 1400Plus transmission electron microscope at 90 kV.

### Nanoparticle tracking analysis

(NTA). NTA of fungal EVs was performed on a LM10 Nanoparticle Analysis System, coupled with a 488 nm laser and equipped with a _S_CMOS camera and a syringe pump (Malvern Panalytical, Malvern, UK) as recently described for cryptococcal EVs (34). All samples were diluted 1:50 in filtered PBS and measured within the optimal dilution range of 9 × 10^7^ to 2.9 × 10^9^ particles/ml. Samples were injected using a syringe pump speed of 50 and three videos of 60 s were captured per sample, with the camera level set to 15, gain set to 3 and viscosity set to water (0.954–0.955 cP). For data analysis, the gain was set to 10 and detection threshold was set to 5 for all samples. Levels of blur and max jump distance were automatically set. The data was acquired and analyzed using the NTA 3.0 Software (Malvern Panalytical).

### RNA isolation and analysis

Vesicular RNA was obtained as previously described by our group (13, 34) with the mirCURYTM RNA isolation kit (Qiagen), used according to the manufacturer’s instructions. As a control we performed RNA isolation from the solid medium alone, which gave negative results (data not shown). For quantitative determination, RNA samples were analyzed with an RNA Agilent 2100 Bioanalyzer (Agilent Technologies) set up for detection of small RNA (sRNA) molecules, as described in recent studies by our group (13, 34). Comparisons between wild-type and mutant cells demanded normalization to the number of vesicles in each sample.

### Analysis of extracellular GXM

The presence of GXM in supernatant fractions was analyzed by ELISA as previously described (35). For GXM quantification in EV fractions, aliquots of 7.8 × 10^8^ EV particles were vacuum dried and disrupted by the addition of 100 μl of a mixture of chloroform and methanol (1:2, v/v). Precipitated polysaccharides were obtained by pulse centrifugation and subsequently delipidated by similar rounds of precipitation using other mixtures of chloroform and methanol (2:1 and 9:1, 100 μl each). The dried precipitates were suspended in PBS (50 μl) for GXM quantification by ELISA (35). Comparisons between wild-type and mutant cells demanded normalization to the number of vesicles in each sample.

### Incorporation of GXM into the surface of acapsular cells

Acapsular *C. neoformans* cells (*cap67*Δ strain) cells were grown in YPD for 24 h, at 30°C with shaking (200 rpm). Yeast cells (5 × 10^6^ cells) were collected by centrifugation and washed twice in PBS. The cells were then suspended in 150 μl PBS containing GXM precipitated from EV samples (10 μg/ml) or 8 × 10^8^ EVs (particle number estimated by NTA) and incubated at room temperature for 24 h. After incubation, the cells were extensively washed with PBS and processed for immunofluorescence as previously described by our group (36). In these assays, the cell wall was stained in blue with calcofluor White and capsular structures appeared in red, after incubation with mAb 18B7. The cells were visualized on a Leica TCS SP5 confocal microscope or analyzed with a FACS Canto II flow cytometer. Data was processed with the FACSDiva Version software, version 6.1.3.

### Scanning electron microscopy

Fungal cells were grown on solid YPD as described before in this section (capsule repression conditions) or incubated in RPMI for capsule induction. For capsule enlargement, 2 × 10^6^ cells were suspended in 200 μl of RPMI and incubated for 24 h at 37°C, under a 5% CO_2_ atmosphere. Cryptococcal cells were washed three times with PBS and fixed in 2.5% glutaraldehyde in 0.1 M sodium cacodylate buffer (pH 7.2) for 1 h at room temperature. The cells were then washed three times with 0.1 M sodium cacodylate buffer (pH 7.2) containing 0.2 M sucrose and 2 mM MgCl_2_. Washed cells were adhered to coverslips that were previously coated with 0.01% poly-L-lysine (Sigma-Aldrich) for 20 min. Adhered cells were gradually dehydrated in ethanol (30, 50 and 70% for 5 min, then 95% and 100% twice for 10 min). The samples were critical point dried (Leica EM CPD300) immediately after dehydration, mounted on metallic bases (stubs), coated with a gold layer of 15-20 nm particles (Leica EM ACE200) and finally visualized in a scanning electron microscope (JEOL JSM-6010 PLUS/LA) operating at 20 kV. For analysis of capsular dimensions, at least 50 cells were analyzed individually and had their total area determined using the ImageJ software (National Institutes of Health, USA). Since no differences in cell bodies were observed between the different strains and conditions used in this study, we assumed that differences in the total cellular area reflected alterations in capsular dimensions.

### Statistics

Statistical analyses were performed with the GraphPad software (La Jolla, CA, USA). Group comparisons were submitted to One-Way ANOVA followed by the Tukey’s multiple comparisons test.

## Acknowledgements

We thank Dr Arturo Casadevall for providing the antibody to GXM (mAb 18B7). We are grateful to Tabata Klimeck (Fiocruz) for preparation of TEM samples, Bruna Marcon (Fiocruz) for help with fluorescence microscopy and Patricia F. Herckert (Fiocruz) for preparation of fungal cultures. We are also thankful to Guilhem Janbon and Frederique Moyrand (Pasteur Institute, Paris) for discussions on the new protocol and for sharing results reproduced in their laboratory. We are grateful to Josh Nosanchuk (Albert Einstein College of Medicice, USA) for helpful suggestions. MLR is currently on leave from the position of Associate Professor at the Microbiology Institute of the Federal University of Rio de Janeiro, Brazil.

